# Identifying Upper Airway and Evaluating Adenoid in Lateral Cephalometric Radiographs of Pediatric Patients Using Image-based Deep Learning Technique

**DOI:** 10.1101/2021.04.19.440369

**Authors:** Kan Yao, Yilun Xie, Wenwen Yu, Silong Wei, Liang Xia, Tong Zheng, Xiaofeng Lu

**Affiliations:** Department of Oral and Cranio-maxillofacial Surgery, Shanghai Ninth People’s Hospital, College of Stomatology, Shanghai Jiao Tong University School of Medicine; National Clinical Research Center for Oral Diseases; Shanghai Key Laboratory of Stomatology and Shanghai Research Institute of Stomatology; Department of Stomatology,Renji Hospital Affiliated to Shanghai Jiaotong University School of Medicine

**Keywords:** obstructive sleep apnea, adenoid hypertrophy, upper airway, image-based deep learning

## Abstract

**Purpose:** Adenotonsillar hypertrophy is considered as one of the primary causes of pediatric obstructive sleep apnea. In clinical practice, pediatric patients with potential obstructive apnea should undergo assessment of upper airway obstruction and adenotonsillar size. As lack of well-trained physicians in China, large numbers of such patients could not be accurately evaluated in time. We attempted to find a better way to assess upper airway obstruction.

**Methods:** We developed a computational method which may allow clinicians to trace upper airway obstruction. We utilized a Detectron2 Mask R-CNN architecture that was pretrained on ImageNet-1k dataset. We then trained it on our COCO dataset using transfer learning for segmentations of upper airway and related key structures. With the instance segmentations, we utilized a tracing algorithm developed by our own to sketch the contours of key segments. At the end, our system would use the traced landmarks to calculate the targeted clinical data.

**Results:** We validated the effective-ness of the algorithm in two steps. First, we used a validation set (COCO dataset) to evaluate the segmentation performance of our system. Our system achieved a mean segmentation AP of 61.30, with airway segmentation AP of 55.64 and cranial base segmentation AP of 66.96. In addition, the AP_50_ was 99.49 and AP_75_ was 73.62. Second,we analyzed the data from our system and experts using a single rater, absolute-agreement, 2-way mixed-effects model, and got ICC value of 0.73 with 95% confident interval = 0.63-0.81.

**Conclusions:** In this study, we created a deep-learning based system to help clinicians evaluate upper airway in lateral cephalometric radiographs which we believe could improve clinical practice. With the evolving of technology, our system would become more integrated into medical care of OSA, freeing the clinical practitioners from repetitive tasks and enabling them to concentrate on improved patient care.

## 1. Introduction

Adenotonsillar hypertrophy is considered as one of the primary causes of pediatric obstructive sleep apnea(OSA,pediatric)[1][2]. The prevalence of adenoid hypertrophy in children was estimated up to 34%[3]. The enlarged adenoid would block the upper airway,resulting sleep apnea and mouthbreathing. Thus, patients with adenoid hypertrophy may suffer from daytime sleepiness,behavior disorders and dento-maxillofacial deformities[1][4][5][6].

In clinical practice, pediatric patients with potential obstructive apnea should should be assessed whether they are sufferd from upper airway obstructions, especially enlarged adenoids and tonsils. Lateral cephalometric radiograph(LCR), a lateral radiograph of the head and neck which is obtained in a standardised cephalostat, is considered as a simple and economical way to diagnose upper airway obstruction[7][8]. It is believed that LCRs are reproducible and comparable among different patients. As a matter of fact, cephalometric analysises based on LCRs are used by orthodontists worldwidely to estimate dento-maxillofacial growth[9][10][11]. In additon, a number of upper airway measurements have been proposed, lots of which aims to identify the obstruction caused by the enlarged adenoid. The widely-accepted methods are McNamara’s line [12]and Fujioka’s adenoid-nasopharyngeal ratio[13]. Both measurements required identification of upper airway and some other related structures in LCRs, and mostly measured by trained physicians. As lack of well-trained physicians in China, large numbers of pediatric patients with potential obstructive apnea could not be accurately evaluated in time. A better way to assess upper airway obstruction becomes a pressing demand.

To settle the problem mentioned above, we developed a computational method which may allow clinicians lacking relevant experience to trace upper airway obstruction. With the rapidly improved object detection and semantic segmentation in the vision community, we are able to build a deep learning system which can automatically assess upper airway. We first utilized a Detectron2 Mask R-CNN architecture[14][15] that was pretrained on ImageNet-1k dataset[16]. The pretrained model was based on a ResNet101+FPN backbone with standard conv and FC heads for mask prediction[17][18]. We then trained it on our COCO dataset[19] using transfer learning[20] for upper airway and related key structures segmentation. With the instance segmentations, we then utilized a tracing algorithm developed by our own to sketch the contours of key segments and find the landmarks. At the end, our system would calculate the targeted clinical data using the traced landmarks. Figure 1 shows the working system. We validated the effectiveness of the algorithm in two steps. First, we used validation set (COCO dataset) to evaluate the semantic segmentation performance of our system. Second, we compared the outcomes of our data with ones of physicians.

**Figure 1:**
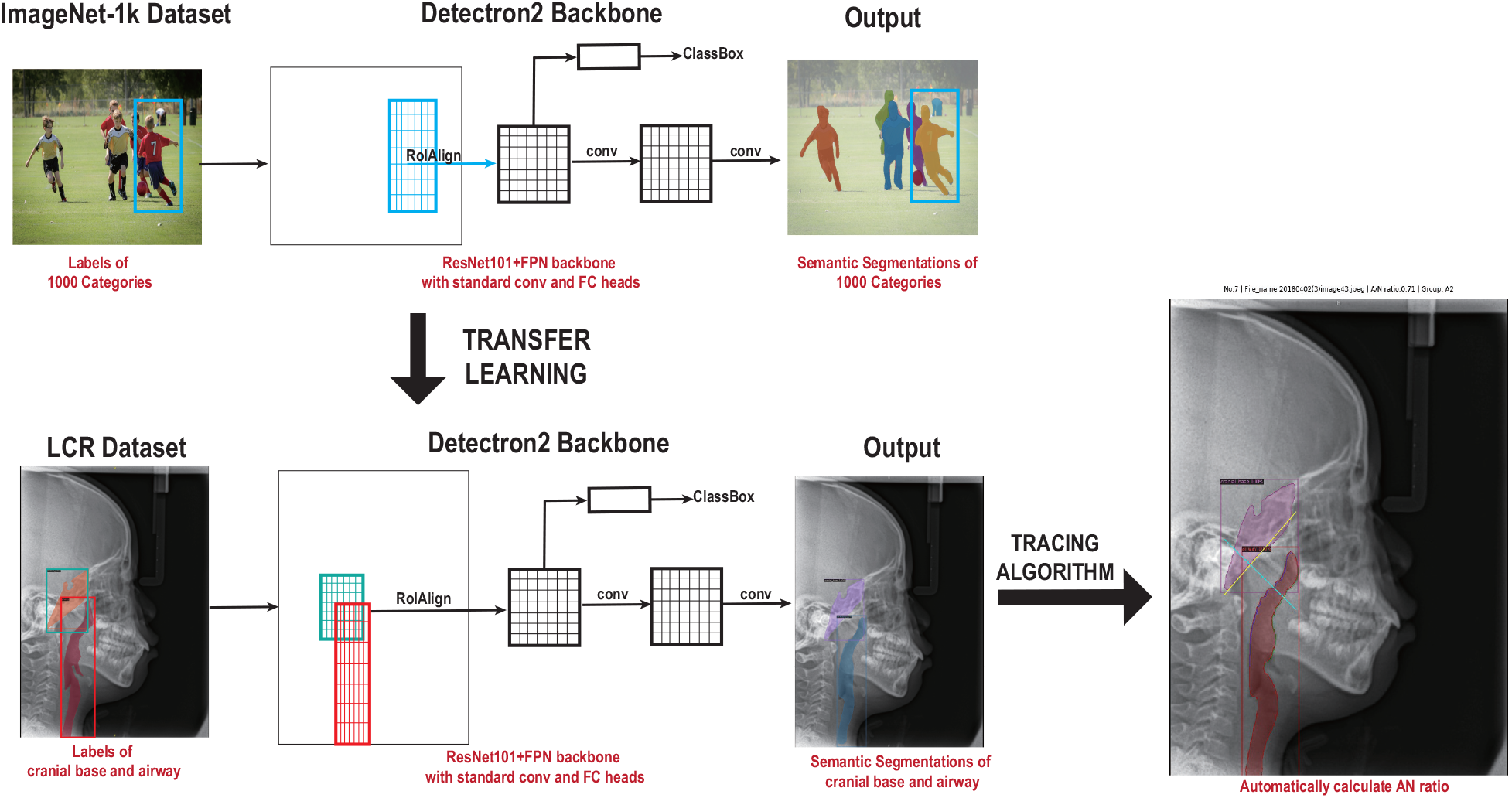
Schematic of the working system.Schematic depicting how a detectron2 trained on the ImageNet dataset of 1,000 categories can be adapted to significantly increase the accuracy and shorten the training duration of a network trained on a novel dataset of LCR images. After segmentations of upper airway and cranial base, our tracing algorithm would automatically figure out the data we need. Note:The deep-learning model is derived from Kaiming He’s paper[14].

## 2. MATERIALS AND METHODS

### 2.1. Datasets

Our data were retrospectively obtained from Department of Oral and Craniomaxillofacial Surgery, Shanghai Ninth People’s Hospital, College of Stomatology, Shanghai JiaoTong University School of Medicine. All the patients came to consulting room with complaints of snoring, mouth breathing or dento-maxillofacial deformities, and the attending physicians decided to take LCR to evaluate upper airway. We randomly picked 510 LCRs which were taken from March 2018 to October 2019 and converted them to JPEG image format. All the patients are anonymized. The LCRs were then labelled by 2 experienced clinicians together using VGG Image Annotator (VIA,version 2.0.8)[21]. In each image, we labelled two key regions, airway and cranial base.(Figure 2). While labeling the airway region, we took the palate and tongue as the anterior wall of airway and ignore tonsils. The annotations were exported to COCO format[19]. We divided the LCRs into two datasets. Four fifth of the data (408 LCRs) were randomly picked as train dataset. The rest of the data (102 LCRs) were considered as validation dataset.

**Figure 2:**
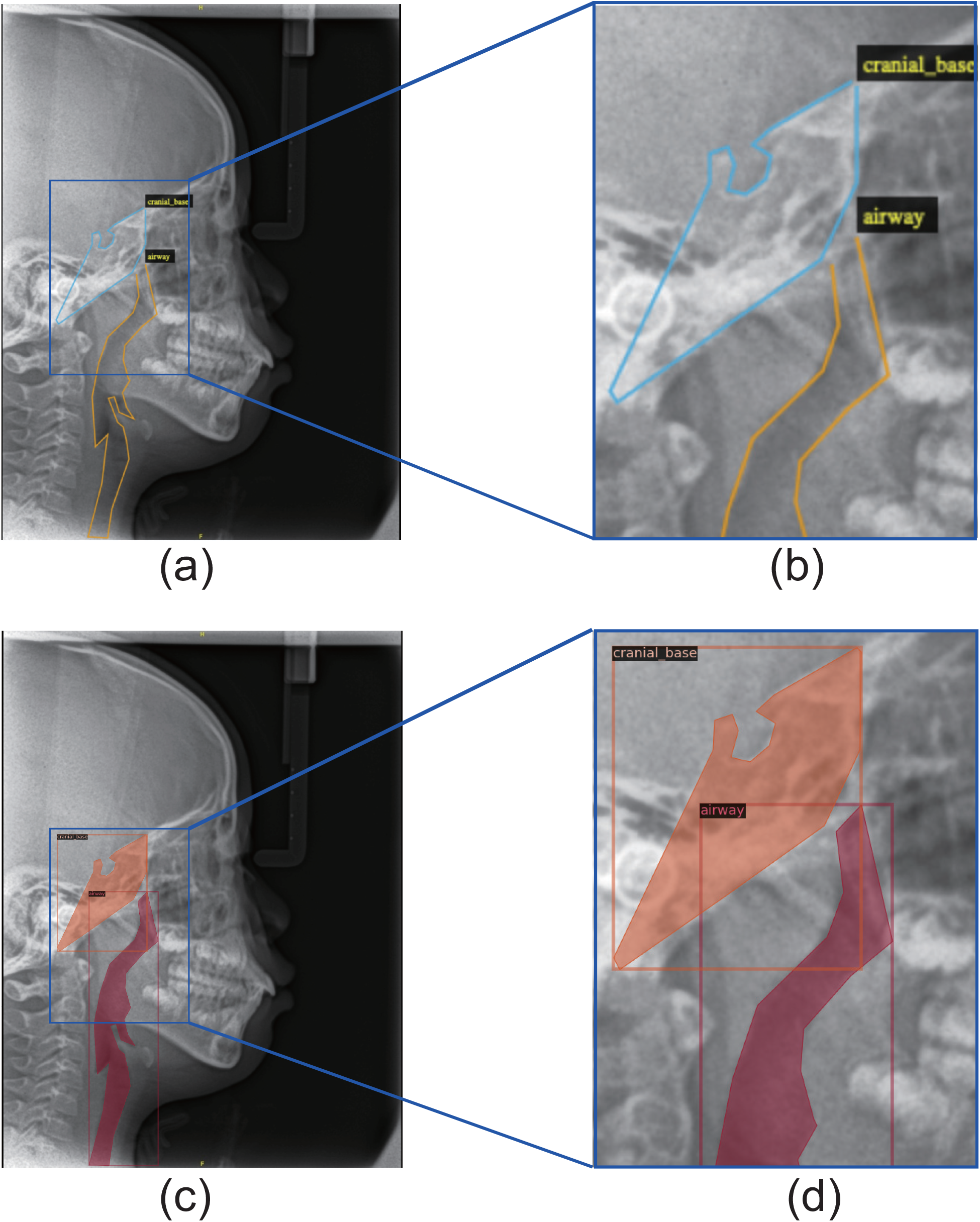
An example of labeling of LCR images.(a) Labeling by experts using VGG Image Annotator (VIA,version 2.0.8); (b) Magnified details of human labeling; (c) Visualize the annotations of sample to make sure the data loading is correct; (d) Magnified details of annotations

### 2.2. Semantic Segmentation Model

We utilized model Detectron2 [15] to train semantic segmentation. Detectron2 is a model that based on Mask R-CNN architecture[14]. Before transfer learning, the model had been pretrained on ImageNet-1k dataset[16] using a ResNet101+FPN backbone with standard conv and FC heads for mask prediction[17][18]. The model was powered by the PyTorch deep learning framework[22]. In this study, we trained our dataset using PyTorch 1.3, CUDA 10.2, cuDNN 7.6.3 with NVIDIA P100 GPU.

### 2.3. Tracing Algorithm

In this study, the tracing algorithm was based on well-performed airway and cranial base segmentation. We took a modified adenoid-nasopharyngeal ratio(AN ratio) as an indicator to test our system’s reliability. The AN ratio was based on Fujioka’s method[13]. The AN ratio we adopted is shown in Figure 3 and is widely-used in China[23]. If a patient’s adenoid is too liitle for us to catch point A’, we regarded the AN ratio as 0. Based on the definition, we designed a module to automatically calculate AN ratio. Firstly, the module would outline the segments. Secondly, it obtained the anterior margin of basiocciput and then carried out the fitting equation of line B through linear regression based on least square method. Thirdly, it seperated the contours of airway into two parts: upper and lower parts. The upper part referred to adenoid margin while the rest reffered to palate margin. Fourthly, the module would try to locate the point A’, N’ and B’ with our algorithm. Then our system would get AN ratio by dividing A(distance between A’ and B’) with N(distance between N’ and B’)(Figure 4).

**Figure 3:**
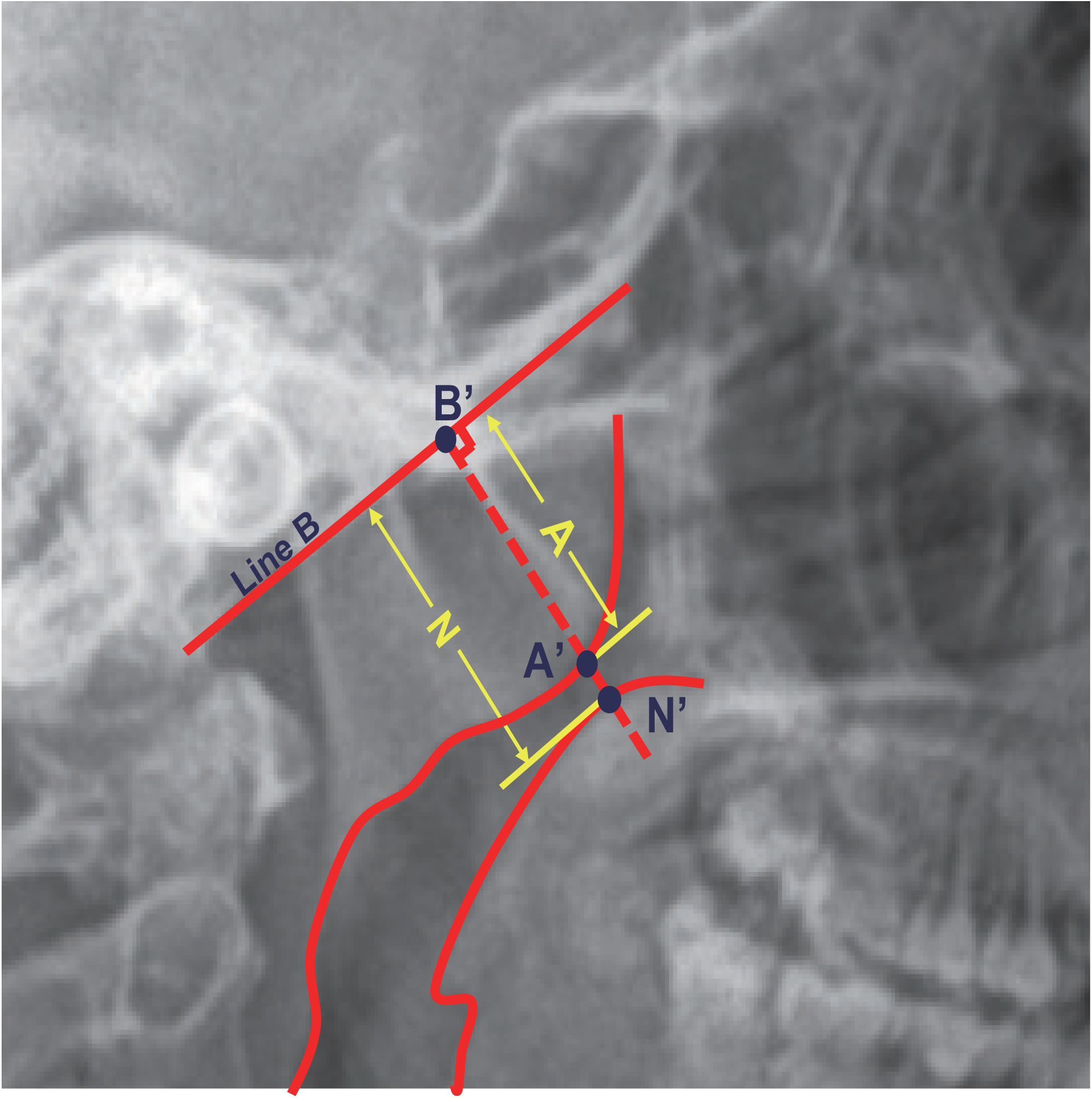
Modified adenoid-nasopharyngeal ratio measurements. “Line B” is the line drawn along straight part of anterior margin of basiocciput. Adenoidal width(A) represents distance from A’, point of maximal convexity, to line B. “A” is measured along line perpendicular from point A’ to its intersection with B, B’ in this figure. Point N’ is the intersection of line A’B’ and the posterior margin of plalate. Nasopharyngeal width(N) is the distance between point B’ and N’. Adenoid-nasopharyngeal ratio(AN ratio) is calculated by dividing A with N.

**Figure 4:**
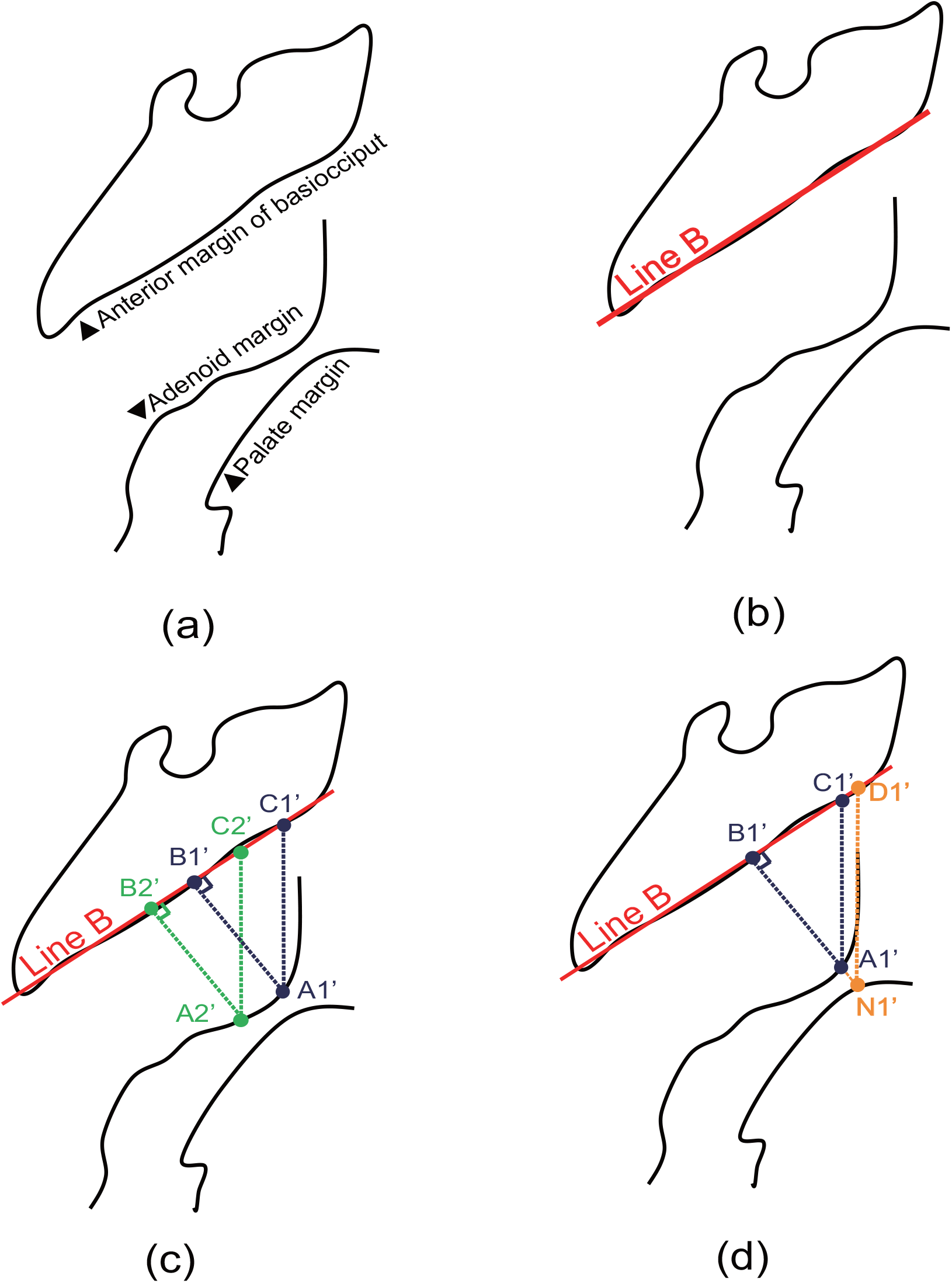
Schematic of tracing alogrithm. Schematic depicting how our tracing alogrithm works. (a)Contours from segmentations of detectron2; (b) Figuring out the line B through linear regression based on least square method; (c) Figuring out the point A’. Ponit C’ is the point on line B which share the same abscissa value with point A’. Distance between A’ and C’ is used to locate the point A’.The point A’ which matches the max distance between A’ and C’ is also the one matches the max distance between A’ and B’; (d) Figuring out the point N’. The point N’ is the intersection of line A’B’ and the plalate margin. Ponit D’ is the point on line B which share the same abscissa value with point N’. AN ratio is calculated by dividing length of line A’C’ with length of line N’D’, which is equal to dividing length of line A’B’ with length of line N’B’.

### 2.4. Validation

We validated the effectiveness of the algorithm in 2 steps. First, we performed a comprasion of Detectron2 segmentation to our labeling on validation the COCO dataset mentioned above. We used the standard COCO metrics including AP (Average Precision at IoU=0.50:0.05:0.95), AP_50_(Average Precision at IoU=0.50), AP_75_(Average Precision at IoU=0.75). AP was evaluating using mask IoU. Second, we compared the data our system exported with the ones physicians measured. Other 2 experienced medical practitioners manually measured AN ratios of the 102 LCRs in validation dataset seperately. Either of the experts completed the tasks twice with one week interval. We took the mean values of the four-time measurements as the final results from experts, which we considered as “golden standard” in this study. Intra-class coefficient correlation(ICC) as shown in Formula 1 was used to test interrater reliability between our system with experts[24].

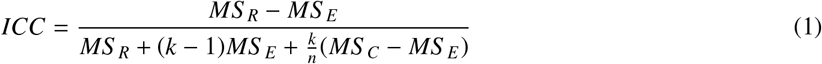

In the formula above, ICC represents intraclass correlation coefficients. MS_*R*_,MS_*E*_ and MS_*C*_represents mean square for rows, mean square for error and mean square for columns, n and k represents number of subjects and number of raters. ICC estimates and their 95% confident intervals were calculated using R version 3.6.3 with irr package (The R Foundation for Statistical Computing, http://www.R-project.org)[25] based on a single rater, absolute-agreement, 2-way mixed-effects model. Values less than 0.50 are indicative of poor reliability, values between 0.50 and 0.75 indicate moderate reliability, values between 0.75 and 0.90 indicate good reliability, and values greater than 0.90 indicate excellent reliability.

## 3. RESULTS

With 4000-Step training, Detectron2 achieved an mean segmentation AP of 61.30, with airway segmentation AP of 55.64 and cranial base segmentation AP of 66.96. In addition, the AP_50_ was 99.49 and AP_75_ was 73.62(Table 1). The detectron2 outputs are visualized in Figure 4. Detectron2 achieved good results in identifying airway and cranial base areas. Based on the semantic segmentation results, the system automatically sketched the point A’, N’ and B’ and figured out the AN ratio(Figure4).

**Table 1:** Instance segmentation AP on COCO test-dev

| AP | $AP_{airway}$ | $AP_{cranialbase}$ | $AP_{50}$ | $AP_{75}$ |
| --- | --- | --- | --- | --- |
| 61.30 | 55.64 | 66.96 | 99.49 | 73.62 |

We analyzed the data from our system and experts using a single rater, absolute-agreement, 2-way mixed-effects model, and got ICC value of 0.73 with 95% confident interval = 0.63-0.81. Thus, we considered the test-reliability of our system as “moderate” to “good”.

## 4. DISCUSSION

### 4.1. Clinical Application

In this study, we described an AI platform for automated evaluation of upper airway, which we believe could assist clinicians dealing with patients with OSA. By employing a transfer learning algorithm, our system showed good performance of upper airway analysis without the need for a highly specialized deep learning machine and without a database of millions of example images. In additon, the performance of our system got “moderate” to “good” test-reliability with clinical experts. Thus, we believed our system was capable of screening out patients with potential upper airway obstructions in LCRs, especially in areas without enough experienced physicians. In additon, our system was designed to deal with images of different formats, which we hope our system would deal with just photographs of X-ray outcomes. We think such a platform could help patients as much as possible, with the fast development of mobile internet. One of our next work is to improve our system’s segmentation performance and to make our system more reliable, especially in photographs of X-rays.

### 4.2. Limitations and Future Work

In our system, we still saw some limitations. We screened out the results from our system and attempted to figure out why system performed poorly in some images. We attributed 2 poor outcomes to poor object segmentations and found 15 outcomes could improve by algorithm optimization in the future (Figure 5). The core of our system is the object segmentation of key areas. Thus, the performance of object segmentation somehow determines the total performance of our system. In the outcomes mentioned above, the poor performance of object segmentations made our system failed to get the A’s and N’s. In the 15 outcomes, we found our tracing alogrithm did not do well when dealing with little adenoids. Our system seemed to be confused about scoring AN ratio as 0. It would figure out the point A’ in LCRs with little adenoids, instead of regarding them as 0. It is worth mentioning that the situation those images our system carried out low values of AN ratios is still acceptable in clinical practice. To solve the problem mentioned above, we plan to add a classifier in our system, which would detect LCRs with little adenoids before calculation.

**Figure 5:**
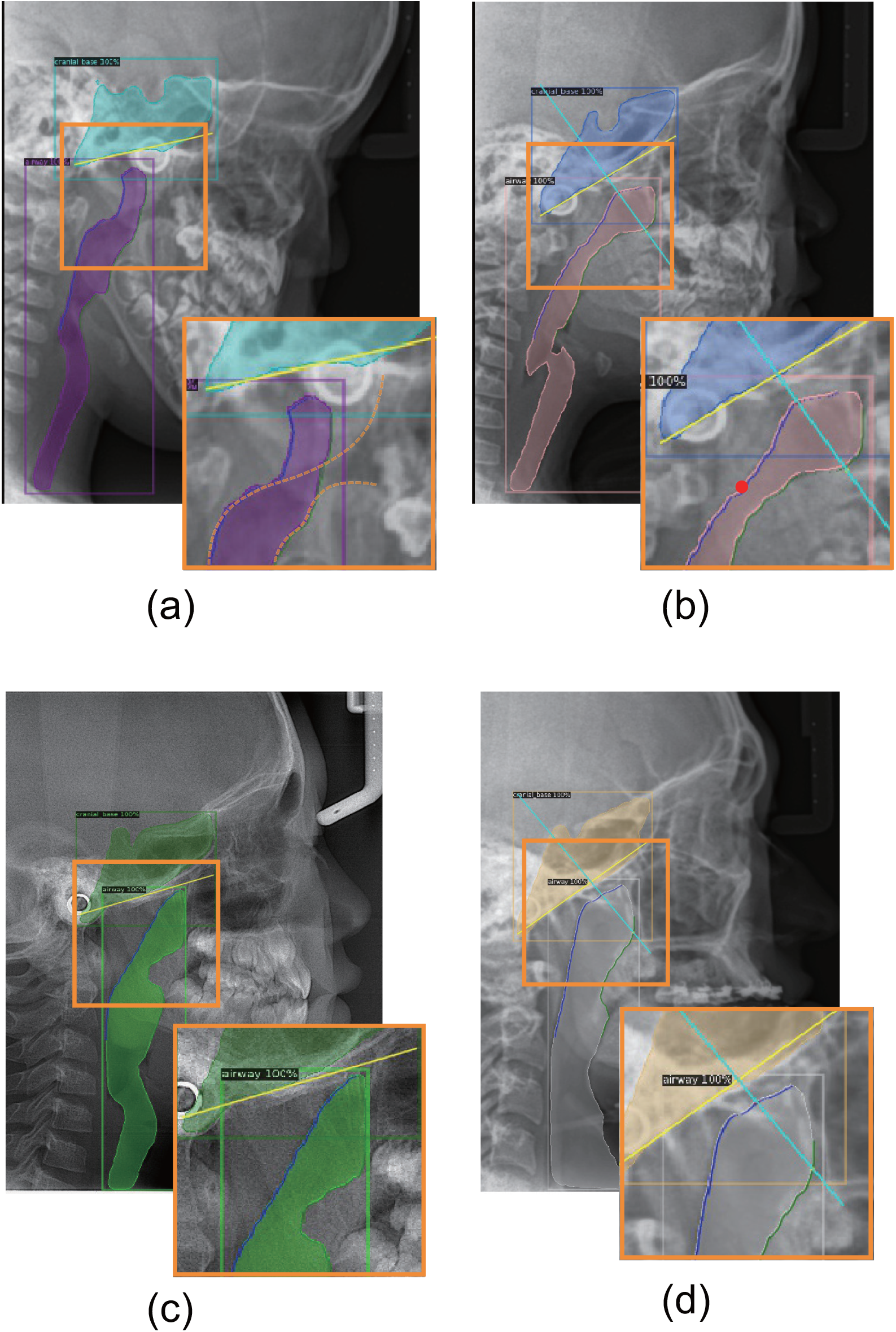
Examples of performance. (a)Poor segmentation of airway.The dashed lines shows the expected airway margins; (b) Failed to get the expected point A’. The red circle represents the expected point A’; (c) Failed to get the point A’. In this case, the adenoid is enlarged, but our algorithm failed to get the point A’; (d) Failed to detect little adenoid. In this case, the AN ratio should be scored as 0, but our system sketched the point A’ and calculated AN ratio.

In the other hand, our work was dealing with 2-dimentional(2D) medical images, which may undergo information loss. In LCRs we could only get anteroposterior and suprainferior diameters of upper airway. As upper airway is a 3-dimensional(3D) structure, the ideal evaluation should consist of anteroposterior, mediolateral and suprainferior diameters. In clincal practice, we could adopt Computerized Tomography (CT) and Magnetic Resonance Imaging(MRI) to obtain 3D airway details. Howerver, higher dimensional data means more amount of information and more complicated structure. As we are new to the area of deep learning and 2-dimentional(2D) cephalometric analysises are still widely and routinely used for upper airway evaluation in China, we chose to design our system capable of dealing with LCR first. Our future work aims to make our system able to treat 3D data. We plan to obtain 3D structures of upper airway in 2 ways. First, we would try CBCT with little radiation. The system would be able to analyse Digital Imaging and Communications in Medicine (DICOM) data from CBCT[26]. Second, we would use endoscopic technique or endosonography to reach 3D structure of upper airway. We hope our system would not only rival human experts in the future, but also dig the hidden data of the relation between OSA and upper airway, which would make clinical practices more effective and accurate.

## 5. CONCLUSION

Image-based deep learning is an emerging tool with huge potential, proper use of which can bring impressive benefits to patients and clinicians. In this study, we created a deep-learning based system to help clinicians evaluate upper airway in LCRs which we believe could improve clinical practice. With the evolving of technology, our system would become more integrated into medical care of OSA, freeing the clinical practitioners from repetitive tasks and enabling them to concentrate on improved patient care.

